# Nucleus reuniens gates selective hippocampal-prefrontal interactions during NREM sleep

**DOI:** 10.64898/2026.11.11.765971

**Authors:** Shan Shao, Jose Ferrero Lopez, Takeshi Kawano, Chinmayi Bhagvat, Dion Khodagholy, Jennifer Gelinas

**Affiliations:** Pediatrics Division of Neurology, University of California Irvine, Irvine, CA, USA; Toyohashi University of Technology, Toyohashi, Aichi, Japan; Department of Biomedical Engineering, University of California Irvine, Irvine, CA, USA; Department of Electrical Engineering and Computer Science, University of California Irvine, Irvine, CA, USA

## Abstract

Memory consolidation requires functional synchrony between the hippocampus (HC) and the medial prefrontal cortex (mPFC). The nucleus reuniens (RE) is a structural and functional hub between HC and mPFC. Here, we used *in vivo* electrophysiology to monitor the hippocampal-thalamic-prefrontal network in rats during non-rapid eye movement (NREM) sleep. We identified ripple-like activity in the RE (120 Hz peak frequency, 45 ms oscillatory duration) that occurred most frequently during NREM sleep and was associated with robust recruitment of thalamic neural spiking. RE ripple activity was phase-locked to both hippocampal sharp wave-ripples (SWR) and mPFC ripples. Although ripples were independently detected in all three structures, we found that coupling between HC and mPFC ripples mainly involved co-occurring RE ripples. These results suggest RE ripples help organize when hippocampal activity engages prefrontal circuits during sleep, with potential implications for network mechanisms of memory consolidation.

## Introduction

Hippocampus-dependent memories are initially encoded through coordinated activity in hippocampal and cortical networks during wakefulness (1–3). During subsequent non-rapid eye movement (NREM) sleep, repeated reactivation of hippocampal representations is thought to support their integration into distributed neocortical networks(4,5). This progressive stabilization and reorganization from labile memory traces into more stable and long-lasting representations is referred to as memory consolidation.

Brain oscillatory activity during sleep facilitates communication between the hippocampus (HC) and neocortical regions, such as medial prefrontal cortex (mPFC) (6). During non-rapid eye movement (NREM) sleep, the neural spiking sequences that represent waking experience are replayed in a temporally condensed manner during hippocampal sharp wave-ripples (SWRs) (7–9). SWRs are critical for memory(10,11) and their interaction with cortical oscillations and neural spiking patterns facilitates network-level consolidation (7,12). Ripple-band activity in cortical regions similarly supports memory processes. These cortical ripples facilitate cross-cortical neuronal interactions (13–15), couple with hippocampal SWRs in a learning-and memory phase-dependent manner (16,17) and are a key feature of neural interactions supporting human memory (18,19).

The medial prefrontal cortex (mPFC) is a key structure implicated in hippocampal-dependent memory consolidation. There is critical memory-related temporal coupling of SWRs with spindles in this region (5,20), but SWR-cortical ripple coupling is also prominent. Independent, compared to co-occurring, SWRs and cortical ripples appear to play different roles in communication between the structures, potentially enabling coordinated integration of recent and remote information (8). The mechanisms governing this temporally selective, gated coupling remain unresolved.

The thalamic nucleus reuniens (RE) is uniquely positioned to regulate these interactions through its reciprocal connections with both the mPFC and the hippocampal formation (21). In addition, the mPFC receives direct, monosynaptic input from hippocampal CA1 via a pathway that exhibits bidirectional synaptic plasticity but does not project back to CA1 (22–24). Together, these anatomically distinct pathways create multiple, yet specific and constrained, routes for information flow through the network.

Correspondingly, RE has been identified as a coordinator of hippocampal-prefrontal activity. RE beta activity precedes hippocampal-prefrontal beta synchronization during memory retrieval, and optogenetic RE activation induces this synchrony (25). Synchronized hippocampal-prefrontal gamma bursts, as well as activation of cell assemblies, occur during slow oscillations and RE inactivation disrupts this coordination under anesthesia (26). Lesions of the RE result in impaired memory persistence(27), indicating a critical functional role.

Here, we use *in vivo* electrophysiology in freely moving rats to simultaneously monitor hippocampus, mPFC and RE activity during NREM sleep. We identify activity patterns in the RE that possess features similar to SWRs and cortical ripples. These RE ripples are associated with robust changes in RE neural spiking and demonstrate significant temporal coupling relationships with both SWR and mPFC ripples. Critically, RE ripples appear to robustly influence the temporal synchronization of SWR with mPFC ripples, such that the probability of a coupled SWR-mPFC ripple is exceedingly low in the absence of RE ripple co-occurrence. Taken together, these results suggest that RE functions as a gate for cross-structure ripple synchronization, potentially enabling directed information flow required for specific memory processes.

## Results

### RE displays ripple-like activity prominently during NREM sleep

To monitor the local field potential as well as the single-unit activity in the RE during sleep and wakefulness, we performed *in vivo* electrophysiological recordings in freely moving rats implanted with 32-channel silicon probes in the RE (Figure 1 A). During NREM sleep, we observed characteristic state-dependent brain oscillations, including thalamocortical sleep spindles and delta waves, on channels histologically confirmed to localize to RE (Supplementary figure 1 A, marked in shade). Unexpectedly, there were also bursts of increased power in the ripple band (100-200 Hz) in these channels, corresponding to recurring high frequency oscillatory patterns in the LFP (Figure 1 B, marked in shade).The power of such ripple-like oscillations was highest in channels localized to the center of the RE and tapered off toward the dorsal or the ventral RE border (Figure 1 C, left).

**Figure 1:**
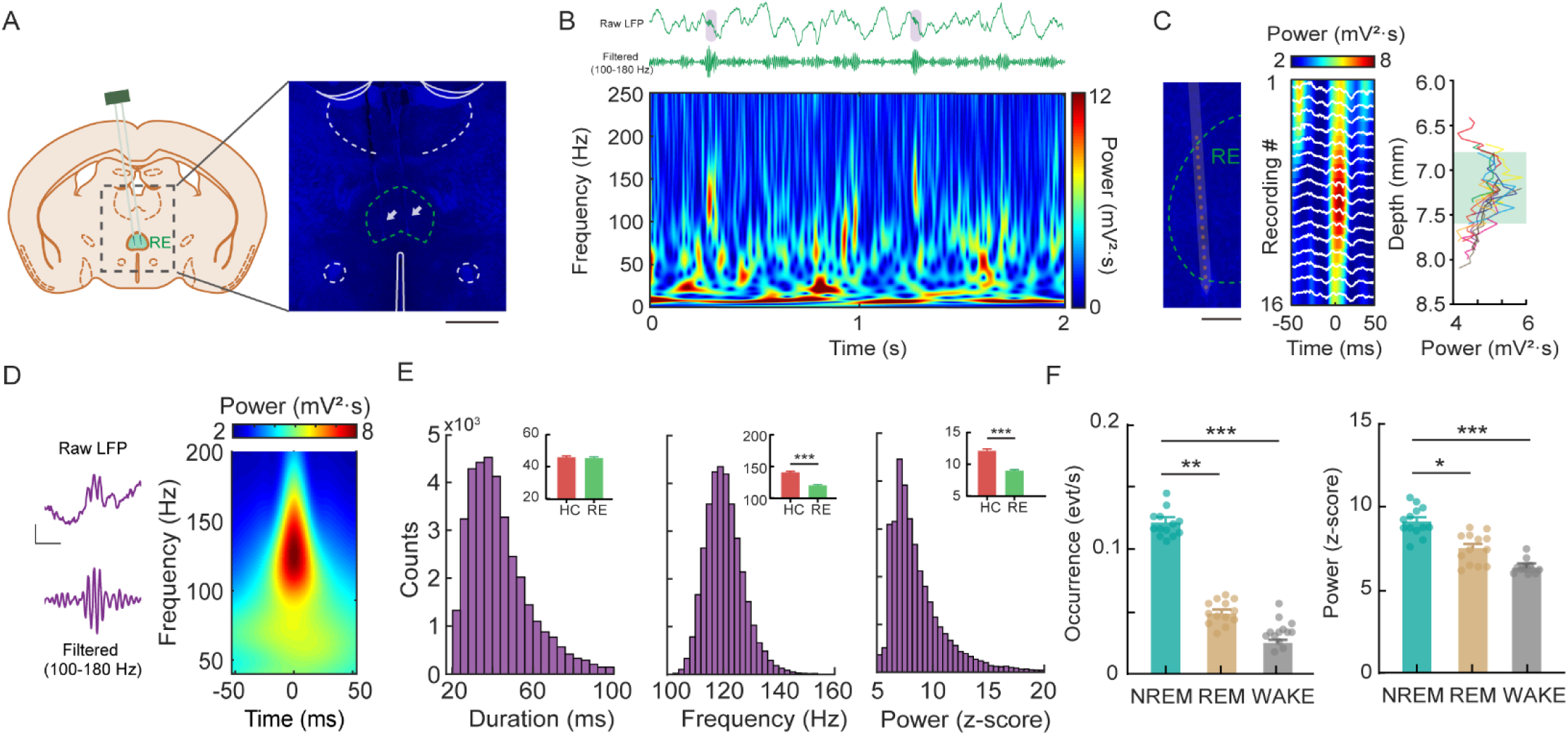
Ripple-like oscillations in RE during NREM sleep. A) Schematic of silicon probe implantation into the RE (left). Representative probe tracks in the RE indicated by white arrowheads (right); scale bar 1 mm. B) Representative wide-band and filtered traces (upper) and sample spectrogram (lower) of RE activity during NREM sleep. C) Sample depth profile of RE ripples recorded along a linear silicon probe shank (16 sites); scale bar 200 um. Sample recording sites from one shank went across the RE (left). Spectrogram of the sample ripple-like activity across the recording channels (middle). Average ripple power along the depth of recording sites across RE (green shaded box indicates RE location relative to brain surface (from -6.8 mm to -7.6 mm)) (n = 175 sites of 11 shanks from 13 rats) (right). D) Sample raw LFP trace and band-pass filtered LFP trace of RE ripples (left); scale bar 0.5 mV, 20 ms. Average RE ripple time-frequency spectrogram (n = 1000 ripples). E) Histograms of RE ripple duration (left), peak frequency (middle) and power (right) (n = 35060 events from 13 rats). Inset demonstrates average characteristics of HC ripples and RE ripples (Wilcoxon rank-sum test, *** p = 0.0002) (n = 13 rats. Error bars indicate ± SEM). F) State-dependent occurrence (left) and power (right) of RE ripples (Kruskal-Wallis test, Occurrence, ** p = 0.0032, *** p = 2.77 × 10^-7^, Power, * p = 0.016, *** p = 1.28 × 10^-6^) (n = 13 rats. Error bars indicate ± SEM).

Similarly, ripple power averaged across rats peaked at dorsoventral coordinates corresponding to the RE (Figure 1 C, right). Because RE has been involved in gamma burst regulation, we quantified gamma-band (50-80 Hz) power over the same time window (± 50 ms from ripple peak). Unlike ripple-band power, gamma-band power did not exhibit a RE-centered dorsoventral gradient (Supplementary Figure 1 B), indicating that this spatial profile is specific to ripple-band activity. Thus, RE appears to exhibit anatomically confined ripple oscillations.

Peak frequency, duration, and power of RE ripples were unimodally distributed (mean peak frequency = 120 ± 8 Hz; mean duration = 45 ±18 ms; mean normalized power = 8.2 ± 2.8 (z-score) (n = 35060 events; Figure 1D-E). Compared to previously well-characterized ripple band activity patterns in hippocampus (sharp wave-ripples), RE ripples were slightly lower frequency and power, but had a similar duration (Figure 1E). When compared to cortical ripples, RE ripples showed shorter duration, slower frequency but higher power (Supplementary Figure 1 C). Across behavioral states, occurrence and power of RE ripples were highest in NREM sleep (Figure 1F), though oscillations with similar duration and frequency were detectable in REM and waking at significantly lower rates (Supplementary Figure 1D). Together, these findings indicated that RE ripples resemble ripples of other brain regions, with preferential occurrence in NREM sleep.

### RE ripples modulate thalamic neural spiking

We next asked whether RE ripples were associated with RE neuronal activity. The majority of recorded RE neurons were significantly modulated around RE ripple-like activity (n = 189/258 neurons), with 48% showing increased firing and 25% showing decreased firing (Figures 2A-B). The proportion of ripple-modulated neurons did not differ significantly along the anteroposterior axis of the RE (n = 11 rats; Supplementary Figure 2B), suggesting that ripple-associated neuronal recruitment was not related to a specific anteroposterior subregion of RE. Baseline firing rates and spike waveform properties differentiated response to RE ripples. RE neurons that increased their firing during RE ripples exhibited increased baseline firing rates and wider action potential waveforms, whereas those that decreased their firing during RE ripples had lower baseline firing rates with increased burstiness (Figure 2C, Supplementary Figure 2C-E). Among neurons with increased firing to RE ripples, 46% demonstrated significant phase-locking (Figure 2D, n = 57/125 neurons). These results demonstrate that RE neurons are robustly modulated during RE ripples, with bidirectional firing responses suggesting engagement of physiologically distinct neuronal populations.

**Figure 2:**
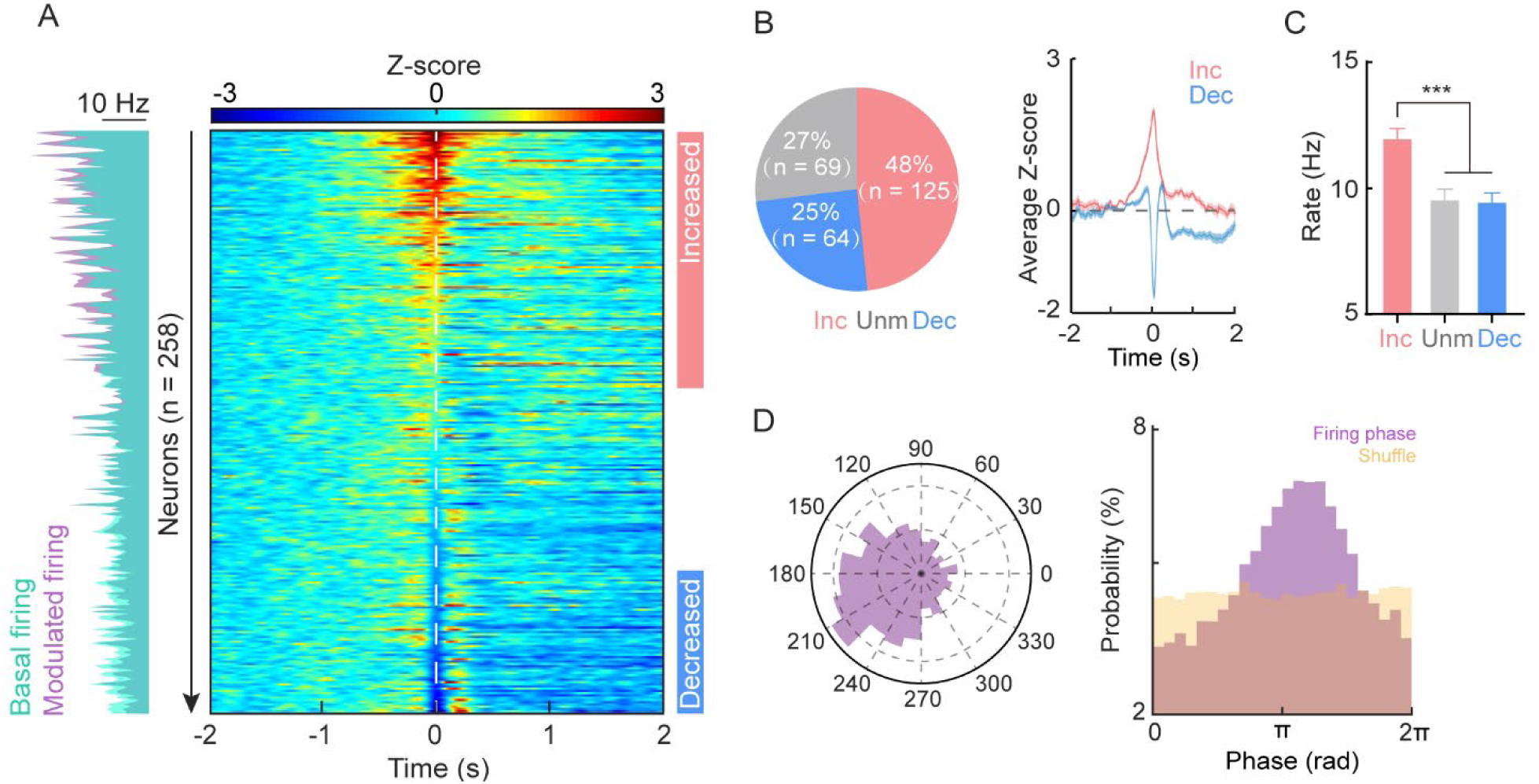
Modulated thalamic neural spiking during RE ripple-like oscillations. A) Mean firing rate of individual neurons before (cyan) and during (purple) RE ripples (left). Heatmap of normalized RE neuron firing centered on RE ripple occurrence (right; colored bars group significantly modulated neurons (ZETA, p < 0.01)). B) Pie chart of proportion of modulated RE neurons; 125 neurons showed increased firing, 64 neurons showed decreased firing and 69 neurons were not significantly modulated (unmodulated) out of all recorded 258 neurons (left). Averaged normalized firing rates of modulated RE neurons during RE ripples (right) ( n= 125 neurons (increased) and 64 neurons (decreased); shades indicate ± SEM). C) Basal firing of different groups of neurons (Kruskal-Wallis test, inc vs unm, ***, p = 9.99 × 10^-6^; inc vs dec, ***, p = 6.20 × 10^-6^, dec vs unm, ns, p = 0.9697, error bars indicate ± SEM). D) Polar plot of a sample phase locking of RE neuron to RE ripples (left). Phase preference of RE neurons relative to RE ripples with shuffling control (right; mean direction = 3.53, concentration factor = 0.27, p = 0.093, n = 57 neurons with significant phase preference).

### RE ripples are coupled with hippocampal and cortical ripples

We next asked how RE ripples interact with hippocampus (HC) and mPFC. We simultaneously recorded activity from the HC, RE, and mPFC during NREM sleep in our rats. We observed that HC and mPFC ripples commonly occurred in close temporal proximity to RE ripples (Figure 3A, Supplementary Figure 3A). Cross-correlation of these activity patterns revealed significant, precise temporal coupling between both HC and RE ripples, as well as mPFC and RE ripples (Figure 3B; Supplementary Figure 3A). RE ripples were significantly phase-locked to HC ripples (distribution centered at 171 degrees), and also to mPFC ripples (distribution centered at 5 degrees; Figure 3C, Supplementary Figure 3B), demonstrating precise oscillatory coordination between all these structures.

**Figure 3:**
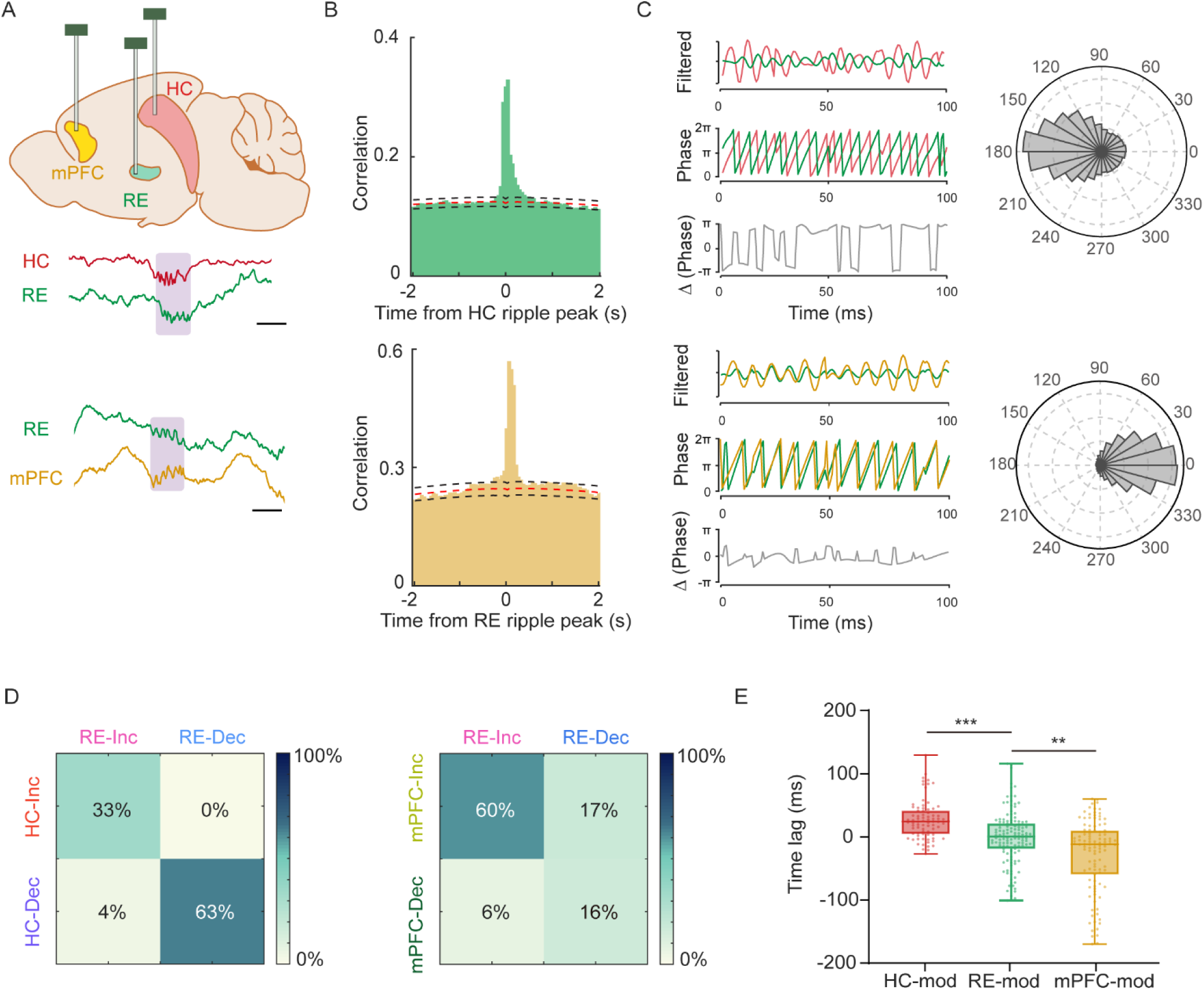
Hippocampal and prefrontal coupling of RE ripples. A) Schematic of device implantation into the HC, RE, and mPFC (upper). Sample raw traces of co-occurring ripples in HC, mPFC, and RE (shaded boxes highlight ripples); scale bar 50 ms. B) Cross-correlogram between HC ripples and RE ripples (upper), RE ripples and mPFC ripples (lower) during NREM sleep (red-dotted line indicates midpoint of upper and lower boundaries of 95% CIs shown as black-dotted lines) (n = 20066 HC ripples, 8466 RE ripples and 16124 mPFC ripples from 13 rats). C) Phase coherence between HC and RE ripples (upper), RE and mPFC ripple (lower). Sample filtered traces, phases and phase differences of co-occurring ripples (left). Polar histogram of phase differences (right; HC vs RE, mean direction = 2.99, concentration factor = 0.63, p < 0.05; RE vs mPFC, mean direction = 0.09, concentration factor = 1.68, p < 0.05). D) Proportion of co-modulated RE neurons by HC and RE ripples (left; n = 70 neurons). Proportion of co-modulated RE neurons by RE and mPFC ripples (right; n = 75 neurons). E) Latencies between co-modulated RE firing peak and ripple peak (Wilcoxon rank-sum test, ** p = 0.0017, *** p = 5.86 × 10^-8^, n = HC-modulated 80 neurons, RE-modulated 189 neurons and mPFC modulated 107 neurons. Error bars indicate Min/Max).

We next examined how RE neurons were modulated during these pairwise ripple interactions. Among RE neurons that were modulated by both HC and RE ripples, 44 of 70 neurons (63%) exhibited decreased firing. However, 45 of 75 RE co-modulated neurons (60%) modulated by both RE and mPFC ripples exhibited increased firing (Figure 3D and Supplementary Figures 3C-D). These opposing response profiles suggest that HC- and mPFC-associated ripple synchrony differentially engage local RE neuronal populations. To examine the temporal organization of these modulations, we quantified the latency between each regional ripple peak and the corresponding peak change in RE firing. HC ripple-associated modulation peaked 25 ms earlier than RE ripple-associated modulation, whereas mPFC ripple-associated modulation peaked 10 ms later (Figure 3E). Together, these results suggest that RE ripples significantly interact with both HC and mPFC ripples, but with distinct neuronal patterns.

### HC-mPFC ripple coupling preferentially involved co-occurrence of RE ripples

Cortical ripples temporally coupled with hippocampal sharp-wave ripples during NREM sleep are implicated in memory consolidation. We investigated how RE is involved in co-occurring HC and mPFC ripples (Figure 4A). Consistent with previous studies, mPFC ripples were significantly temporally coupled to HC ripples (Supplementary Figure 4A). We first separated mPFC ripples based on their temporal coupling with RE ripples. mPFC ripples that had a co-occurring RE ripple were strongly coupled to HC ripples, whereas those without concurrent RE ripple activity had minimal to no significant coupling to HC ripples (Figure 4B-C). Thus, RE ripples appear to robustly gate the temporal synchronization of HC with mPFC ripples. Although the correlation of HC- and RE-ripples was stronger when mPFC ripples were additionally coupled, HC-ripples maintained a significant correlation with RE ripples regardless of mPFC activity (Supplementary Figure 4B-C). Ripples that engaged all 3 structures simultaneously (HC-RE-mPFC) had longer cortical duration compared those that did not, potentially indicating an enhanced response to this synchronized activity pattern (Supplementary Figure 4D).

**Figure 4:**
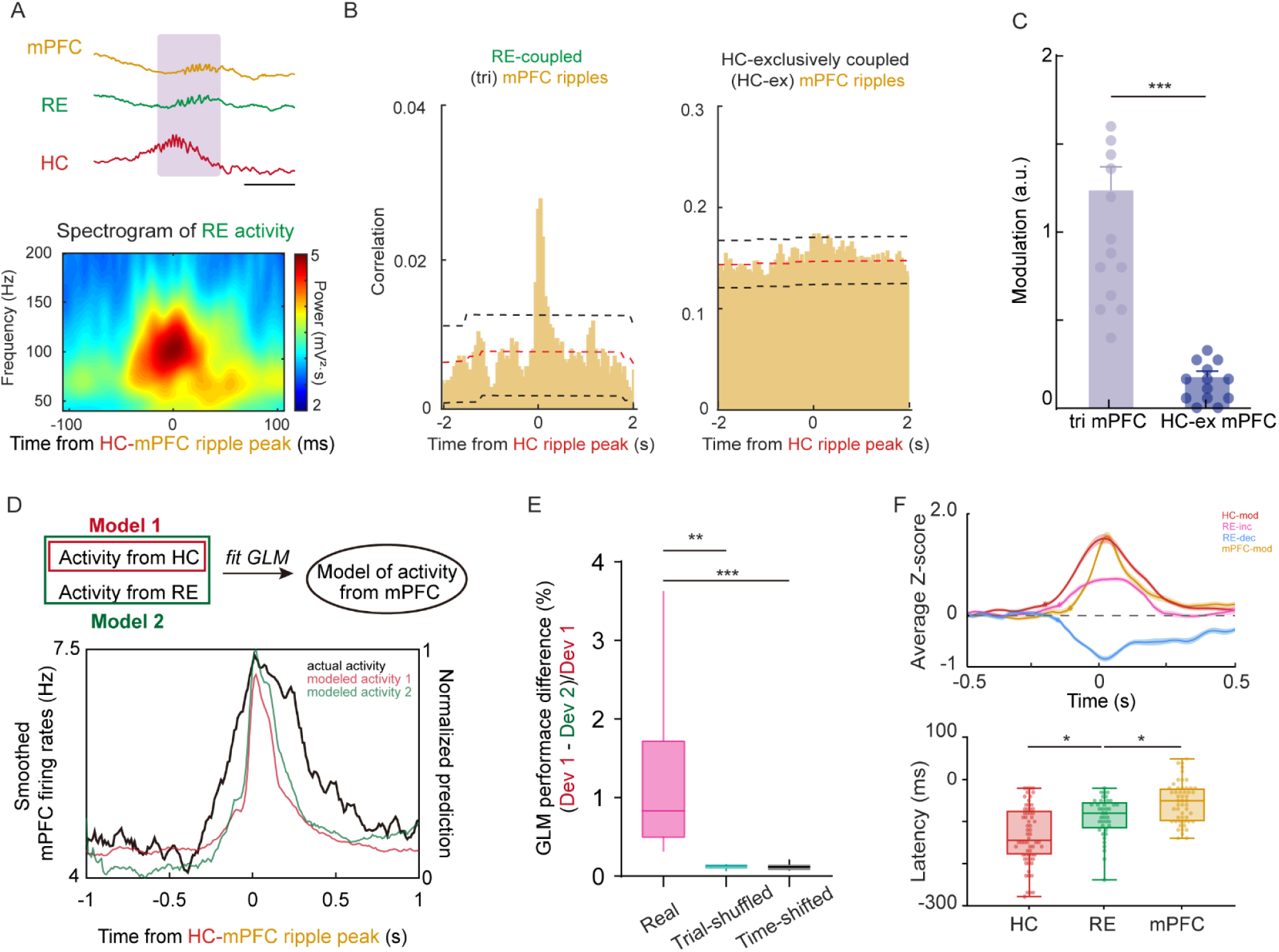
RE ripples gate HC-mPFC ripple coupling. A) Sample raw traces of co-occurring ripples in HC, RE and mPFC (upper; shaded box highlights ripples. Sample spectrogram of RE activity during HC-coupled mPFC ripples (lower). B) Cross-correlogram between HC ripples and RE-coupled mPFC ripples (i.e. ripples that occur across all 3 structures; n = 20066 HC ripples and 1799 mPFC ripples from 13 rats; red-dotted line indicates midpoint of upper and lower boundaries of 95% CIs shown as black-dotted lines) (left). Cross-correlogram between HC ripples and HC-exclusively coupled (HC-ex) mPFC ripples (i.e. ripples that occur only in HC and mPFC, not RE; n = 20066 HC ripples and 14470 mPFC ripples from 13 rats; red-dotted line indicates midpoint of upper and lower boundaries of 95% CIs shown as black-dotted lines) (right). C) Quantification of coupling modulation for tri-synchronized mPFC ripples (HC-RE-mPFC) and HC-mPFC only ripples across individual rat (Wilcoxon Signed-Rank test, *** p = 0.0002) (N = 13 rats. Error bars indicate ± SEM). D) Schematic showing GLMs used to model mPFC activity during HC-coupled mPFC ripples in one recording session as a function of simultaneous HC activity (model 1) or as a function of both HC activity and RE activity (model 2; upper). Sample modeled mPFC activity (lower). E) Relative difference in GLM model performance when adding RE activity as an additional predictor, compared to that using RE trial-shuffled and RE time-shifted controls (Wilcoxon Signed-Rank test, ** p = 0.0028, *** p = 0.0002) (n = 11 sessions from 5 rats. Error bars indicate Min/Max). F) Normalized firing of modulated neurons during HC-coupled mPFC ripples (upper; stars indicate first significant deviation from baseline). Quantification of latencies between HC coupled-mPFC ripple peak and first firing deviation of modulated neurons from each region (Wilcoxon rank-sum test, HC vs RE, * p = 0.012, RE vs mPFC, * p=0.0298) (n = 60 HC neurons, 40 RE neurons and 52 mPFC neurons. Error bars indicate Min/Max).

To further assess whether RE activity contributes to HC-mPFC coordination, we constructed Poisson generalized linear models (GLM) predicting mPFC spiking from HC spiking activity alone (model 1) or from the combined activity of HC and RE (model 2) (Figure 4D). Notably, adding temporally aligned RE activity significantly improved model performance (normalized reduction in deviance) compared to control models in which RE activity was trial-shuffled or temporally shifted (Figure 4E). Consistent with the deviance analysis, inclusion of real RE activity also reduced mPFC spiking prediction error (Supplementary Figure 4E). Together, these findings suggest simultaneous RE activity provides predictive information about mPFC neuronal activity during hippocampal-prefrontal communication.

To better understand the temporal organization of this network, we compared the onset of neuronal recruitment across the HC, RE, and mPFC during mPFC ripples that had a concurrent HC ripple. Response onset was defined as the first time point within ±1 s of the mPFC ripple peak at which normalized firing departed from its baseline mean by more than 3 SD. Above threshold neuronal activity was first detected in HC, followed by RE and then mPFC (Figure 4F). This HC-RE-mPFC sequence, consistent with the temporal ordering of RE neuronal modulation by ripples in the three regions (Figure 3E), supports an ordered progression of involvement across the hippocampal-thalamic-prefrontal network during HC ripples. Together, these findings suggest a role of RE in gating ability of HC ripples to influence mPFC.

## Discussion

Here, we examined how the RE participates in ripple coordination between the hippocampus and mPFC during NREM sleep. We identified ripple oscillations in RE that were most prominent during NREM sleep with modulated thalamic firing. Co-occurrence of RE ripples predicted strong synchronization between HC and mPFC. During this hippocampal-prefrontal integration, HC neurons were firstly recruited, followed by RE and then mPFC. Temporally aligned RE spiking also improved predictions of mPFC firing from hippocampal activity. These findings suggest that RE helps shape the coordination of hippocampal SWR to prefrontal circuits during NREM sleep.

During hippocampal SWR, synchronous firing of CA3 provides excitatory input to CA1, producing the sharp wave profile (7). In CA1, local parvalbumin-expressing interneurons provide phase-locked inhibition to pyramidal cells, helping pace the accompanying ripple oscillation (28,29). During cortical ripples, putative pyramidal cells and interneurons fire at preferred phases, with pyramidal cells firing slightly earlier, consistent with feedback inhibition pacing the rhythm (16,30). We also observed that RE neurons mostly increased their firing during RE ripples, with 46% showed significant phase preference to the ripple cycle. Given the scarcity of local GABAergic interneurons in rodent RE (31), a candidate source of inhibition is the thalamic reticular nucleus (TRN), whose anteromedial sector projects to RE (32). Recruitment of this pathway could transiently suppress RE firing and align subsequent spikes when inhibition subsides (33). Further studies combining simultaneous TRN and RE recordings with selective manipulation of TRN inputs to RE could test whether this pathway is involved in RE ripple generation.

RE ripples engaged neurons with distinct intrinsic properties. Neurons that showed increased firing during RE ripples had higher basal firing rates and wider extracellular spike waveforms, whereas neurons that showed decreased firing had lower basal rates and greater burstiness of firing pattern. This diversity is consistent with the range of firing patterns observed in RE neurons in vitro, where burst propensity depends on membrane potential (33). Retrograde tracing also points to an anatomical basis for heterogeneity. Hippocampus-projecting neurons occupy a broad mediolateral span of RE, while mPFC-projecting neurons are enriched along the midline and in the lateral perireuniens (34). Multi-shank recordings with finer spatial resolution across these subregions, combined with identification of projection targets, could reveal whether the firing profiles we observed map onto distinct output pathways and help explain how different subsets of RE neurons are recruited for hippocampal-prefrontal communication.

The significantly stronger hippocampal-prefrontal ripple coupling observed when RE ripples co-occurred suggests that RE activity may help define a brief window in which mPFC neurons are more responsive to hippocampal input. The improved prediction of mPFC firing with temporally aligned RE spiking is consistent with a contribution that depends on timing. An in vitro study showed that RE input recruits both recurrent excitation and feedforward inhibition in mPFC pyramidal neurons (35). These findings suggest that RE may gate the prefrontal response to hippocampal input during network ripple synchronization by temporally recruiting recurrent excitation in mPFC, thereby enabling a brief window in which coordinated hippocampal and prefrontal activity supports memory stabilization during NREM sleep.

Thalamic ripple events have been reported in human intracranial EEG data, where thalamic sampling has recently become part of clinical practice. Identification of these patterns supports the capacity for thalamic nuclei to exhibit such high frequency oscillatory patterns. Weak, longer time-scale coupling between thalamic ripple events and hippocampal or cortical ripples has been reported in this human data (36). Due to sparse sampling of the human thalamus with relatively large electrodes, it is difficult to ascertain whether highly localized ripples similar to those we identify in RE would be reliably detected across human subjects. It is possible that thalamic-mediated ripple synchronization is strongly network specific, such that assessment of anatomically broad interactions obscures the high temporal precision coupling that we identified in rodents.

The coordination between hippocampal and prefrontal ripples facilitates memory updating and stabilization(16,17). Our observed RE-mediated network interaction may contribute to memory persistence by gating the stabilization of prefrontal representations. Lesions of the RE and rhomboid nuclei impaired long-term spatial recall (27) and prevented a delayed increase in dendritic spines in the anterior cingulate cortex after spatial learning (37). In our recordings, tri-coupled mPFC ripples had longer duration compared to uncoupled ripples. These longer events could give hippocampal and prefrontal ensembles more time to fire together during particular phases of the memory process, creating conditions for synaptic changes that stabilize long-term cortical representations.

## Materials and Methods

### Animals

Adult Long Evans rats of both sexes, weighing between 250 and 350 g, were used in the experiments as stated in the text. Rats were maintained on a regular 12-h light/dark cycle and provided ad libitum access to food and water. All experimental procedures were carried out in accordance with the guidelines of the Institutional Animal Care and Use Committee at Columbia University Irving Medical Center.

### Stereotaxic Surgery

A two-shank silicon probe with 16 recording sites per shank (NeuroNexus) and a five-wire tungsten electrode assembly mounted on a custom microdrive were prepared. Rats were anesthetized with an isoflurane/oxygen mixture (2% for induction, 0.75-1% for maintenance) and placed in a stereotaxic apparatus (David Kopf Instruments). A heating pad was placed beneath the rats to maintain body temperature. Midline incisions were made in the scalp using surgical blades and craniotomies were opened using a dental drill. Wire devices were implanted in the left hippocampal CA1 (-3.5 mm anterior-posterior (AP), -3.00 mm medial-lateral (ML) and −2.20 to −2.40 mm dorsal-ventral (DV)) and left mPFC (3.5 mm AP, 0.2 mm ML and -2.50 mm DV). Probes were implanted in reuniens nucleus (RE; -1.45 mm AP, 0.6 mm ML and 6.90 to 7.20 mm DV, 8 degree from the vertical axis, relative to the brain surface), fixed by C&B-Metabond Quick adhesive Luting Cement (Parkell) and dental cement (Pearson Dental). The craniotomies were covered by Gelfoam and sealed using a 10:1 mixture of paraffin and mineral oil. The reference and ground wires implanted over the cerebellum. During all surgeries, animals were injected subcutaneously with Ketoprofen (2 mg/kg), Bupivacaine (2 mg/kg), and Buprenorphine (0.05 mg/kg). For recovery, animals were placed in a clean cage on a heating pad. Animals were allowed to fully recover from surgery for at least 5 days before further experimentation.

### Histology

After completion of the experiments, electrode placement was verified in all rats. Rats were euthanized with sodium pentobarbital and perfused with PBS followed by 4% paraformaldehyde (PFA). Brains were extracted, postfixed in 4% PFA for 24 h, embedded in OCT (Optimal Cutting Temperature) compound, and sectioned coronally at 60 μm using a cryostat (Leica). Sections were permeabilized in PBS containing 0.25% Triton X-100, stained with DAPI (1:10,000) for 20 min, and mounted with Fluoromount-G (Invitrogen). Images were acquired using a fluorescence microscope (Keyence), and tungsten wire or silicon probe tracks were reconstructed across adjacent sections to determine recording locations. Data from recording sites outside the intended regions of interest were excluded from subsequent analyses.

### *In vivo* electrophysiology recording

For at least five days prior to recordings, rats were habituated to experimenter handling and attachment of cables for electrophysiology recordings. During the habituation, hippocampal electrodes were adjusted in the dorsal-ventral axis to span the layers of CA1 based on localizing neurophysiological signals. For RE recording, electrodes were minimally adjusted to maximize acquisition of neural spikes and then maintained at these coordinates throughout the remainder of the recording procedure.

### Data acquisition and analyses

Neural signals were collected at a sampling rate of 20 kHz by a multichannel extracellular recording system (Intan Technologies). Data were amplified by a recording headstage and digitalized by a USB interface board. The recorded data were analyzed using MATLAB (MathWorks) and visualized using Neuroscope (38).

#### Local field potential (LFP) preprocessing

Data were resampled to 1250 Hz to facilitate LFP analysis. Sleep epochs were determined by the absence of movement in the motion signal recorded by a 3-axis accelerometer and the absence of electromyogram (EMG) artifacts. REM and NREM sleep epochs were classified using a validated automated sleep scoring algorithm based on ratios of cortical delta (0.5-4 Hz), cortical gamma (50-80 Hz) and hippocampal theta (5-8 Hz)(39), followed by visual inspection and manual adjustments to ensure the accuracy of sleep scoring.

#### Frequency domain analysis

Time-frequency representations of the LFP were generated using an analytical Gabor wavelet transform. Spectral power was calculated as the squared magnitude of the wavelet coefficients, yielding spectrograms that characterize changes in oscillatory power over time. Ripple-associated spectrograms were aligned to detected ripple peaks to examine the temporal and spectral structure of activity surrounding ripple events.

To characterize the dorsoventral distribution of ripple- and gamma-band power, 11 probe shanks from 13 rats that crossed the RE at similar AP coordinates were selected. At each recording site, power in the ripple (100-160 Hz) and gamma (50-80 Hz) bands was averaged over a 100-ms window centered on RE ripple peaks, allowing comparison of their spatial profiles along the dorsoventral axis.

#### Ripple detection

For hippocampal ripple detection and initial screening for RE ripples, LFP was band-pass filtered from 100 to 200 Hz, then rectified and normalized. Events where the envelope was two standard deviations above the baseline and peaks surpassed five standard deviations were identified. Events were included when the duration exceeded 20 ms and the duration between two events was more than 30 ms. Events common to the ripple channel and the EMG electrode noise channel were excluded. Cortical and RE ripples were detected using the same procedure as hippocampal ripples, with band-pass filtering adjusted to 110-150 Hz and 100-160 Hz, respectively. The same amplitude thresholds, duration and inter-event interval criteria, and EMG-based artifact exclusion were applied across all ripple detections.

#### Single-unit activity processing

Spike sorting was performed on band-pass-filtered data (250-2500 Hz) using KiloSort (v1.0). Clusters were manually curated in Phy (v2.0) to isolate single units. Single units were further validated based on the mean waveform, autocorrelogram, and consistency of waveform localization with probe geometry. Units lacking a clear refractory period in the autocorrelogram were excluded. Event-related firing-rate modulation was assessed for individual neurons using the Zenith of event-based time-locked anomalies (ZETA) approach(40), which evaluates time-dependent changes in firing without requiring predefined analysis bins. For visualization and z-score analyses, firing rates were calculated in 10-ms bins and normalized to the mean and standard deviation of activity during the 1-s baseline period from 2 to 1 s before event onset.

#### Neuronal phase preference

The instantaneous phase of the ripple-band LFP was extracted using the Hilbert transform of the band-pass-filtered signal. The phase at each spike time was assigned to the corresponding spike, and each neuron’s preferred phase was calculated as the circular mean of its spike phases. Phase-locking strength was quantified using the mean resultant length, and departure from a uniform phase distribution was assessed with Rayleigh’s test. Neurons were classified as significantly phase-locked when p < 0.05 and κ > 0.1.

#### K-means clustering

K-means clustering was performed in MATLAB using two extracted waveform features: trough-to-peak latency and spike asymmetry index. Neurons were assigned to one of three clusters based on Euclidean distance to the nearest cluster centroid. Cluster membership was then used to compare the action potential properties of modulated and unmodulated neurons.

#### Cross-correlogram (CCG) calculation

Temporal coupling between ripple events was quantified using cross-correlograms (CCGs) (17). For each pair of regions, ripple events in one region served as reference events aligned to time zero, and events in the other region were counted in successive time bins of 50 ms relative to each reference event. CCGs were constructed from events occurring during NREM sleep. The statistical significance was determined using a modified convolution method (41), each CCG was convolved with a partially hollowed window. Confidence limits were derived from a Poisson distribution whose mean was determined by this convolution. CCG peaks exceeding the upper confidence limit were considered evidence of significant temporal coupling. Coupling strength was quantified as the relative elevation of the CCG peak (a) above the chance level (b):

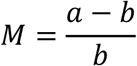

This normalized modulation (M) expresses the excess co-occurrence relative to chance. The same calculation was applied consistently across the conditions included in each comparison.

#### Coupled ripple identification

In the pairwise CCGs, six 50-ms lag bins spanning −150 to +150 ms exceeded the upper bound of the 95% CI. Based on this window, we classified coupled ripple events if their peak times differed by less than 150 ms. mPFC ripples that had co-occurring ripples in both HC and RE were identified in two steps: mPFC ripples were first classified as RE-coupled using the above coupling criterion, and the RE-coupled subset was then similarly classified for HC coupling. HC-mPFC only coupled ripples were identified as events that met the HC-coupling criterion but were not coupled to an RE ripple within 150 ms.

#### Phase coherence

All LFP signals were band-pass filtered using the same 100-200-Hz frequency range before phase extraction to ensure that phase differences were comparable across regions. The instantaneous phase of each signal was then calculated from its Hilbert transform. For each pair of coupled events, the phase difference between the overlapping portions of the two signals were calculated. Phase differences were pooled within each recording session, and phase coherence was quantified using the mean resultant length, and departure from a uniform phase distribution was assessed with Rayleigh’s test.

#### Modulation latency

The normalized firing rates for significantly modulated RE neurons were calculated around ripple events in each region to compare the timing of RE neuronal modulation by ripples in the HC, RE, and mPFC. For each region, the time of each neuron’s peak or trough response was identified. Modulation latency was defined as the interval between this response time and the peak of the corresponding regional ripple. Then these modulation latencies across the three anatomical structures were compared to determine whether RE neuronal modulation followed a temporal order.

#### Generalized linear modeling of HC-RE-mPFC interactions

An adapted generalized linear modeling (GLM) approach (42) was used to determine whether RE activity provides additional information about mPFC activity beyond that explained by hippocampal activity. Analyses were performed separately for each recording session using neural activity aligned to HC-coupled mPFC ripples within a ±0.5-s peri-ripple window. When multiple units were modulated within a region, their activity was averaged within each session. HC and RE activity were used as predictors, whereas mPFC spike counts were used as the response variable in 10-ms bins.

For each session, two Poisson GLMs were fitted. The HC-only model (Model 1) was used to predict mPFC spike counts from HC spiking using a lag kernel spanning −100 to +100 ms relative to each mPFC time bin:

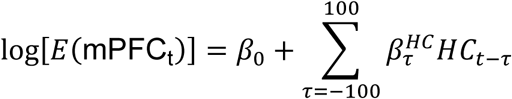

The HC+RE model (Model 2) included the same HC predictors together with RE spiking over the same lag range:

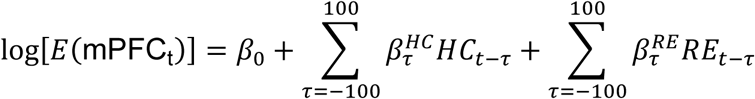

The additional predictive contribution of RE was quantified as the normalized reduction in model deviance:

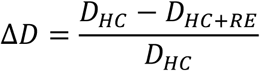

where D_HC_ and D_HC+RE_ represent the residual deviances of Models 1 and 2, respectively. Positive values therefore indicate improved prediction of mPFC activity after inclusion of RE activity.

To determine whether this improvement depended on the actual relationship between RE and mPFC activity, two RE-based shuffle controls were performed while HC activity was kept unchanged. For the RE trial-shuffle control, RE activity was randomly reassigned across events within each session, thereby disrupting the event-specific correspondence between RE and mPFC activity while preserving the temporal structure of RE activity. For the RE temporal-shift control, RE activity within each event was circularly shifted in time, thereby disrupting its temporal alignment with mPFC activity while preserving the temporal structure of the RE activity. Each shuffle procedure was repeated 100 times per session, and the mean shuffled Δ*D* was used for session-level comparisons.

As an additional measure of prediction accuracy, mean squared error (MSE) between observed and predicted mPFC spike counts was calculated within ±50 ms of the ripple peak. MSE was compared across HC-only, HC+RE, trial-shuffled RE, and temporally shifted RE models. All model comparisons and statistical analyses were performed at the session level.

#### Neuronal activation during coupled events

The temporal order of neuronal recruitment across the HC, RE, and mPFC was assessed during HC-coupled mPFC ripple events. Firing activity was aligned to the HC-coupled mPFC ripple peak and normalized to each neuron’s baseline (-2 to -1 second relative to ripple peak) mean and standard deviation. For each significantly modulated neuron, recruitment onset was defined as the first time point within the ±1-s analysis window at which firing exceeded the baseline mean by 3 standard deviations or fell below it by 3 standard deviations.

### Quantification and statistical analysis

All statistical analyses were performed in MATLAB or GraphPad Prism 10. Data are presented as mean ± SEM unless otherwise specified. Paired comparisons were assessed using the Wilcoxon signed-rank test, whereas comparisons between independent groups were assessed using the Wilcoxon rank-sum test. Comparisons across three or more conditions were performed using the Friedman test for repeated measurements and the Kruskal-Wallis test for independent groups. Post hoc pairwise comparisons were performed using Dunn’s multiple-comparisons test. Adjusted p values were used to determine significance for pairwise comparisons. The association between anteroposterior coordinate and the proportion of ripple-modulated neurons was assessed using linear regression. Associations between GLM performance and the number of recorded neurons were assessed using Spearman’s rank correlation. Significant event-related modulation of neuronal firing was identified using the ZETA test at p < 0.01. Neuronal phase locking and interregional phase synchronization were assessed using Rayleigh’s test for departure from a uniform phase distribution at p < 0.05 and κ > 0.1. Significant temporal coupling between ripple events was determined using Poisson-derived 95% confidence limits for cross-correlograms. For other analyses, statistical significance was defined as p < 0.05. Significance levels are indicated as * p < 0.05, ** p < 0.01, and *** p < 0.001.

## Data and code availability

All data needed to evaluate the conclusions are present in the manuscript and/or the Supplementary Information. Any additional data and relevant code are available from the corresponding author upon request.

