## Supplementary Figures for "Nucleus reuniens gates selective hippocampal-prefrontal interactions during NREM sleep"

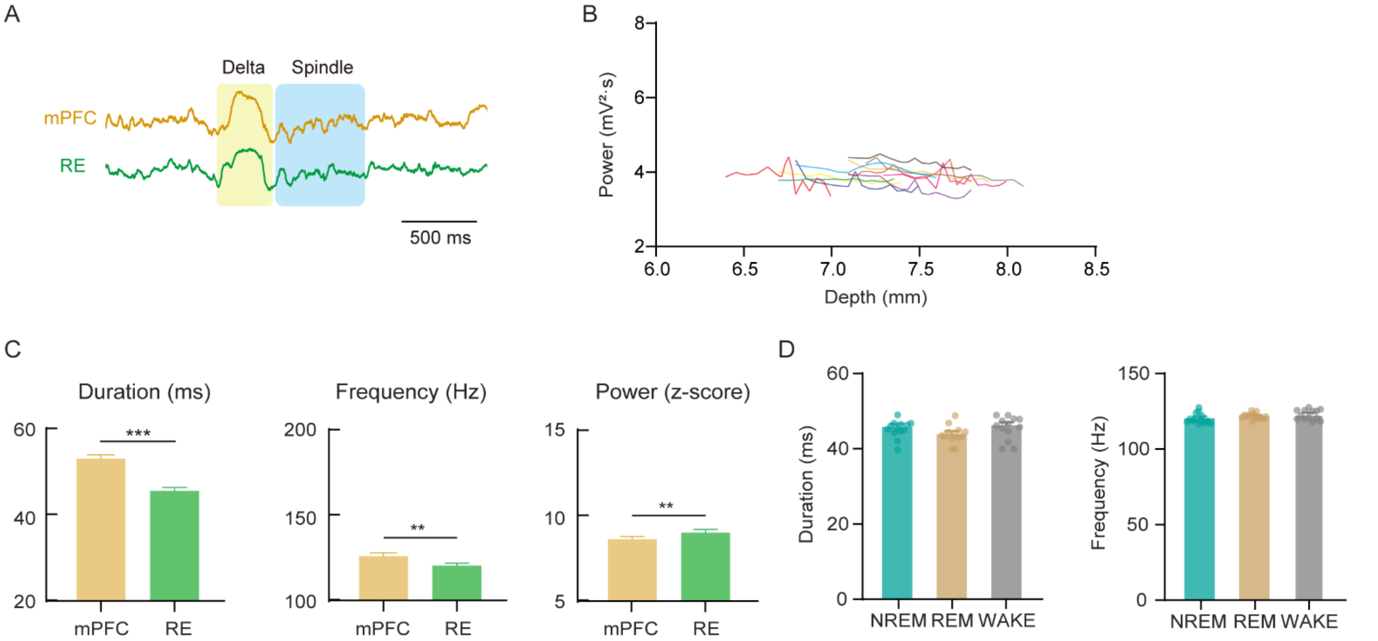


**Supplementary figure 1**

A) Representative wide-band activity of RE and mPFC during NREM sleep.
B) Average gamma power along the depth of recording sites across RE (n = 175 sites of 11 shanks from 13 rats)
C) Average characteristics of mPFC ripples and RE ripples (Wilcoxon rank-sum test, ***, p = 0.0002, **, p = 0.0012, ** p = 0.0024) (n = 13 rats. Error bars indicate ± SEM).
D) Average duration (left) and average peak frequency (right) of RE ripples during NREM, REM and wake state (N = 13 rats. Error bars indicate ± SEM).

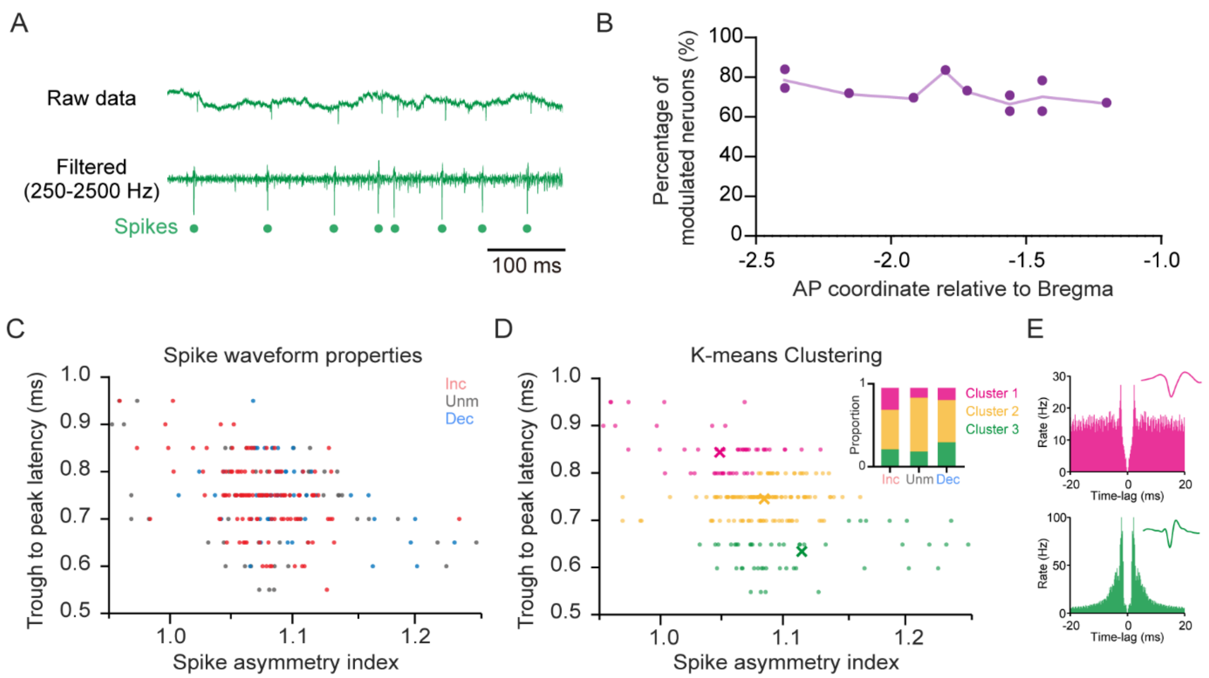


**Supplementary figure 2**

A) Sample wide-band and filtered traces and detected thalamic neural spiking.

B) Proportion of modulated neurons along the anteroposterior (AP) axis of RE (linear regression, p = 0.58, n = 11 sites from 11 rats).

C) Action potential properties of modulated and unmodulated neurons.

D) Proportions of three K-means clusters in modulated and unmodulated neurons based on action potential properties.

E) Auto-correlograms and action potential waveforms of sample neurons from Cluster 1 and 3.

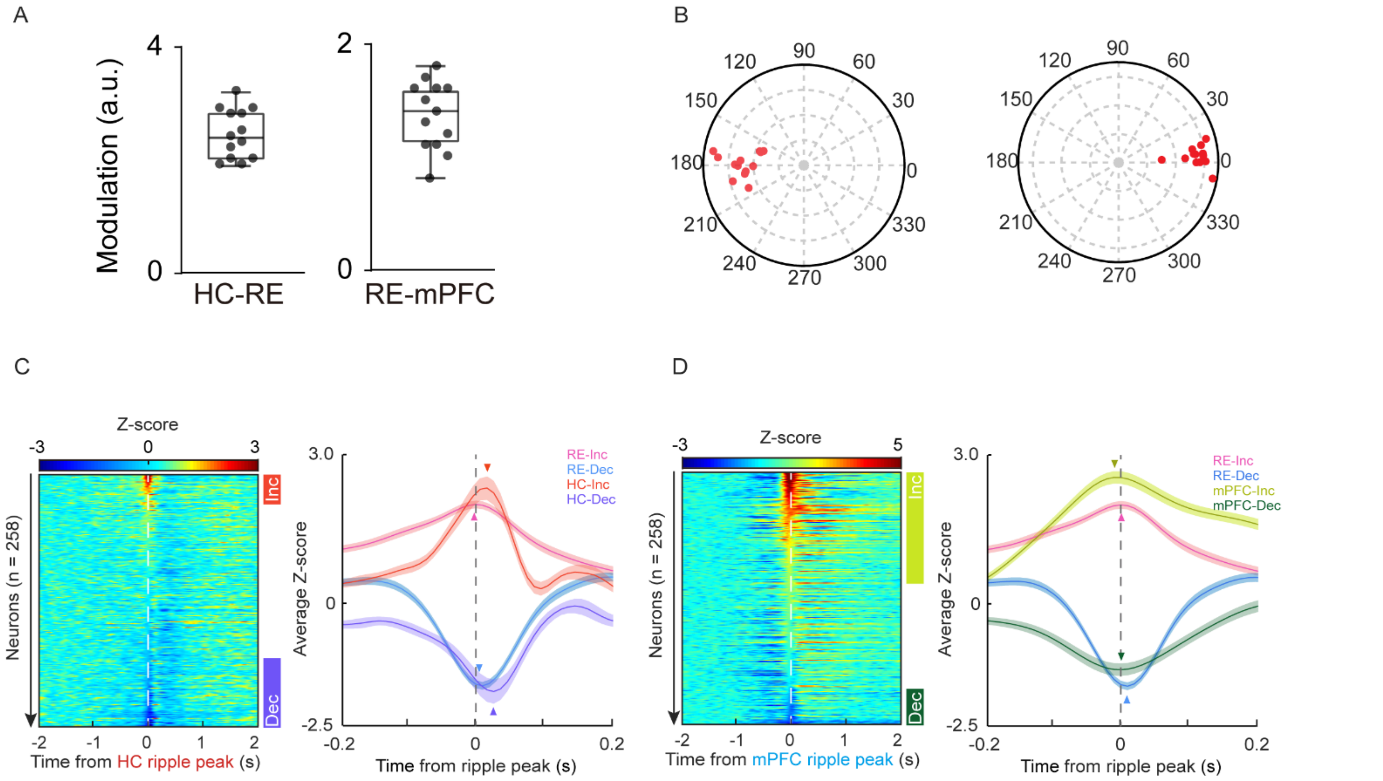


**Supplementary figure 3**

A) Coupling modulation of HC and RE ripples (left) and RE and mPFC ripples (right) from a representative individual rat (N = 13 rats. Error bars indicate Min/Max).

B) Phase coherence values from across individual rats (N = 13 rats).

C) Heatmap of normalized firing of RE neurons during HC ripples (colored bars on the right indicate significantly modulated neurons at $p < 0.01$). Averaged activity of modulated RE neurons during HC and RE ripples (right; n = 80 and 189 neurons. Arrowheads indicate the modulation peak).

D) Heatmap of normalized firing of RE neurons during mPFC ripples (colored bars on the right indicate significantly modulated neurons at $p < 0.01$). Averaged activity of modulated RE neurons during mPFC and RE ripples (right; n = 107 and 189 neurons. Arrowheads indicate the modulation peak).

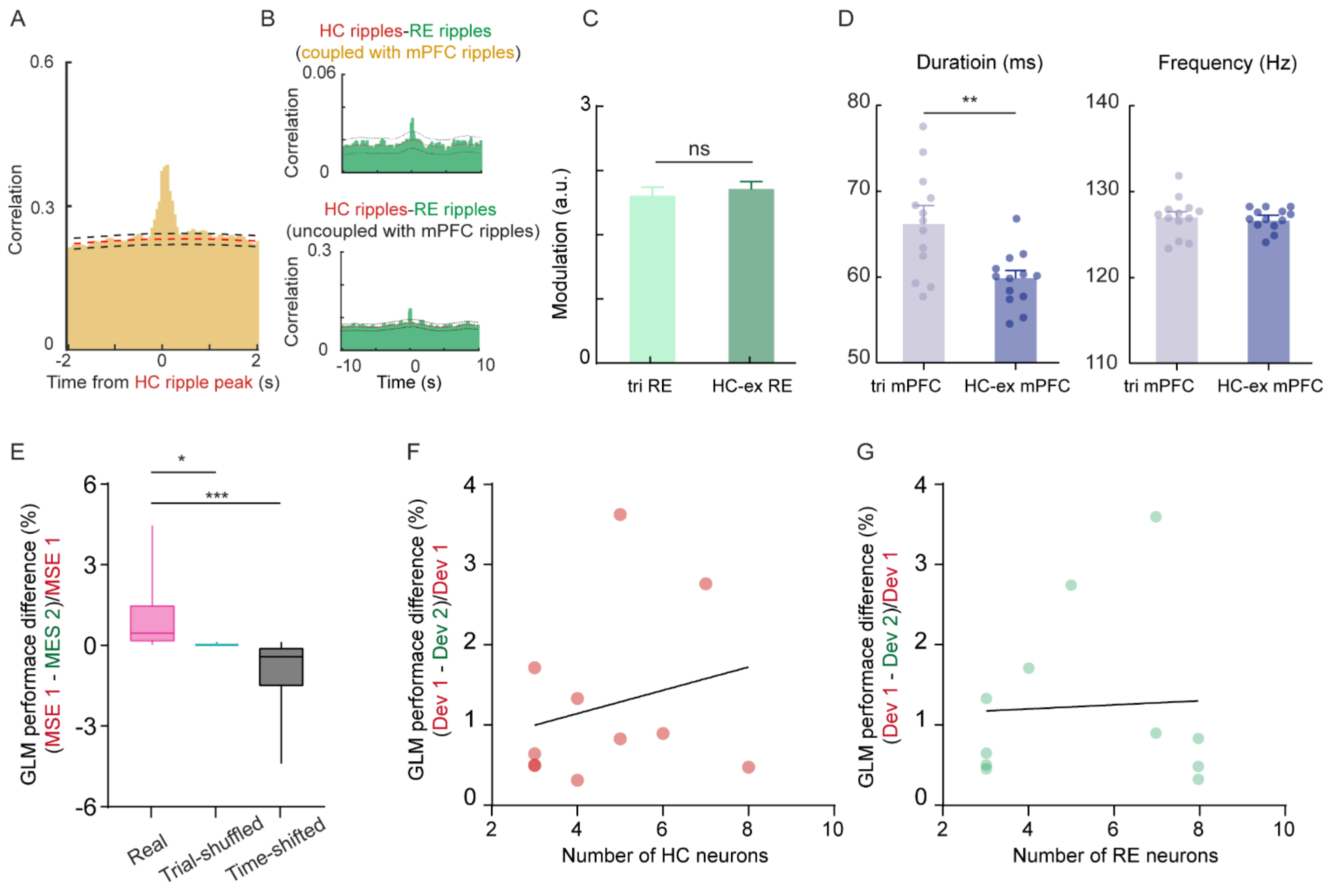


**Supplementary figure 4**

A) Cross-correlogram between HC ripples and mPFC ripples during NREM sleep (n = 20066 HC ripples and 16124 mPFC ripples from 13 rats; red-dotted line indicates midpoint of upper and lower boundaries of 95% CIs shown as black-dotted lines).

B) Cross-correlogram between HC ripples and mPFC-coupled RE ripples (upper; n = 20066 HC ripples and 1799 RE ripples from 13 rats). Cross-correlogram between HC ripples and HC-exclusively coupled RE ripples (lower; n = 20066 HC ripples and 6667 RE ripples from 13 rats) (red-dotted line indicates midpoint of upper and lower boundaries of 95% CIs shown as black-dotted lines).

C) Quantification of coupling modulation for tri-synchronized RE ripples and HC-RE only ripples across individual rat (Wilcoxon Signed-Rank test, $p = 0.52$) (N = 13 rats. Error bars indicate ± SEM).

D) Average characteristics of tri-synchronized and HC-mPFC only ripples (Wilcoxon Signed-Rank test, ** $p = 0.0034$) (N = 13 rats. Error bars indicate ± SEM).

E) GLM model performance difference (MSE: mean square error) (Wilcoxon Signed-Rank test, * $p = 0.038$, *** $p = 0.0001$) (n = 11 sessions from 5 rats. Error bars indicate Min/Max).

F) No significant linear correlation between GLM performance and number of HC neurons (Spearman correlation, $r = 0.17$, $p = 0.61$) (n = 11 sessions from 5 rats).

G) No significant linear correlation between GLM performance and number of RE neurons (Spearman correlation, $r = -0.15$, $p = 0.65$) (n = 11 sessions from 5 rats).
